# Introduction of Kinesin-13 family specific residues increases the microtubule end residence time of a Kinesin-1

**DOI:** 10.1101/405654

**Authors:** Hannah R. Belsham, Claire T. Friel

## Abstract

Kinesins that regulate microtubule dynamics, such as the Kinesin-13 MCAK, require the ability to recognise the microtubule end. All microtubule regulating kinesins studied to date have this ability and thus exhibit increased microtubule end residence times. In contrast, purely translocating kinesins such as Kinesin-1 do not need to recognise the microtubule end for their function. The residues K524, E525 and R528 in the α4 helix of the Kinesin-13, MCAK, are critical for microtubule end recognition. Here, we show that introducing these Kinesin-13 family-specific residues into a Kinesin-1 increases its microtubule-end residence time up to 4-fold. However, this increase in end residence is not sufficient to confer microtubule depolymerisation activity to a Kinesin-1.

**Significance Statement:** The introduction of Kinesin-13 family specific residues from the α4 helix of the microtubule depolymerising kinesin, MCAK, into a Kinesin-1, increases the microtubule-end residence time between 2 and 4-fold. This demonstrates both the significance of these residues in modulating microtubule end residence, and the capacity to tune kinesin function by modifying the microtubule binding face of the motor domain using protein engineering. Increasing the microtubule end residence time in this way is not sufficient to confer depolymerisation activity to a Kinesin-1. Thus, indicating that the ability to recognise and reside at the microtubule end is not the only determinant of depolymerisation activity. The Kinesin-13 motor domain may also possess the ability to actively break interactions between tubulin subunits.

## Introduction

A key feature of the molecular mechanism of kinesins that regulate microtubule dynamics is the ability to distinguish the microtubule end from the microtubule lattice. Both microtubule polymerising and depolymerising kinesins have been shown to possess the ability to recognise the microtubule end. For example the Kinesin-5, Eg5, is a microtubule polymerase which is important in spindle assembly and axon branching and can slide microtubule bundles (Sawin, LeGuellec et al. 1992, Kapitein, Peterman et al. 2005, Freixo, Martinez Delgado et al. 2018). It stabilises microtubule ends, promoting polymerisation and reducing catastrophe and resides at microtubule ends for 7 seconds (Chen and Hancock 2015). The Kinesin-8, Kip-3, is a microtubule depolymerising kinesins that translocases to the plus end of microtubules where it typically removes only a single tubulin dimer (Varga, Leduc et al. 2009). Kip-3 can remain at the microtubule end for tens of seconds (Varga, Leduc et al. 2009). Other examples of kinesins with extended end residence include the Kinesin-7 CENP-E (Sardar, Luczak et al. 2010), the Kinesin-4 Kif4 (Subramanian, Ti et al. 2013), and NOD (Cui, Sproul et al. 2005), all of which are suggested to have the ability to alter microtubule dynamics.

The Kinesin-13 MCAK, a microtubule depolymerase, also has the ability to distinguish the microtubule end from the lattice (Hunter, Caplow et al. 2003, Friel and Howard 2011). Previously, we have shown that the α4 helix of the MCAK motor domain is critical to this ability (Patel, Belsham et al. 2016). Mutation of the Kinesin-13 family-specific residues K524, E525 and R528 significantly impairs the ability of MCAK to recognise the microtubule end. The mutations K524A, E525A and R528A mutations reduce both the microtubule end residence time and the rate constant for microtubule end stimulated ADP dissociation (Patel, Belsham et al. 2016). As these residues are clearly crucial to the ability of MCAK to recognise the microtubule end, we wondered whether introduction of these Kinesin-13 specific residues into the α4 helix of a Kinesin-1 could confer the ability to recognise the microtubule end to a purely translocating kinesin.

Members of the Kinesin-1 family are considered purely translocating kinesins and the classic cargo carriers. They are highly processive and can operate *in vitro* as homodimers with two identical motors domains bound by a C-terminal coiled-coil tail domain moving in a hand-over-hand fashion along microtubules (Howard, Hudspeth et al. 1989, Yildiz, Tomishige et al. 2004). The motor domains are linked to the tail via a short neck domain, which determines directionality (Case, Pierce et al. 1997). The globular C-terminal end of the tail domain is involved in binding of cargo such as vesicles, organelles and mRNA (Hirokawa, Noda et al. 2009). This function requires abilities such as co-ordination of the interaction of the two motor domains with the microtubule, but does not require the ability to distinguish between microtubule end and microtubule lattice. Here, we introduce Kinesin-13 family specific residues from the α4 helix into the corresponding positions in a Kinesin-1 and show that these mutations increase the microtubule end residence time relative to the wild-type.

## Results

### The Kinesin-1 residues G262, N263 and S266 correspond to K524, E525 and R528 in Kinesin-13

The residues K524, E525 and R528 are critical to the ability of the Kinesin-13, MCAK, to distinguish the microtubule end from the microtubule lattice (Patel, Belsham et al. 2016). To investigate the functional malleability of the kinesin motor domain, we wanted to substitute these Kinesin-13 family specific residues into the equivalent positions in a Kinesin-1 motor domain to see if it is possible to confer microtubule end recognition to a purely translocation kinesin. We used the rat kinesin-1 construct rKin430-GFP (Rogers, Weiss et al. 2001) as a representative Kinesin-1. The amino acid sequence of rKin430 and MCAK (Kinesin-13) were aligned using the Huang and Miller algorithm and BLOSUM50 scoring matrix (Huang and Miller 1991), to identify the residues in Kinesin-1 that correspond to the Kinesin-13 residues K524, E525 and R528. By sequence alignment the corresponding Kinesin-1 residues were found to be G262, N263 and S266 (Figure 1A). Analysis of the crystal structures of Kinesin-1 and Kinesin-13 (Figure 1B) confirms the spatial correspondence of these residues which are found on the microtubule binding face of the α4-helix in both Kinesin-1 and Kinesin-13. It is notable that the Kinesin-13 residues have large charged side-chains whilst the corresponding Kinesin-1 residues are smaller and/or uncharged.

**Figure 1:**
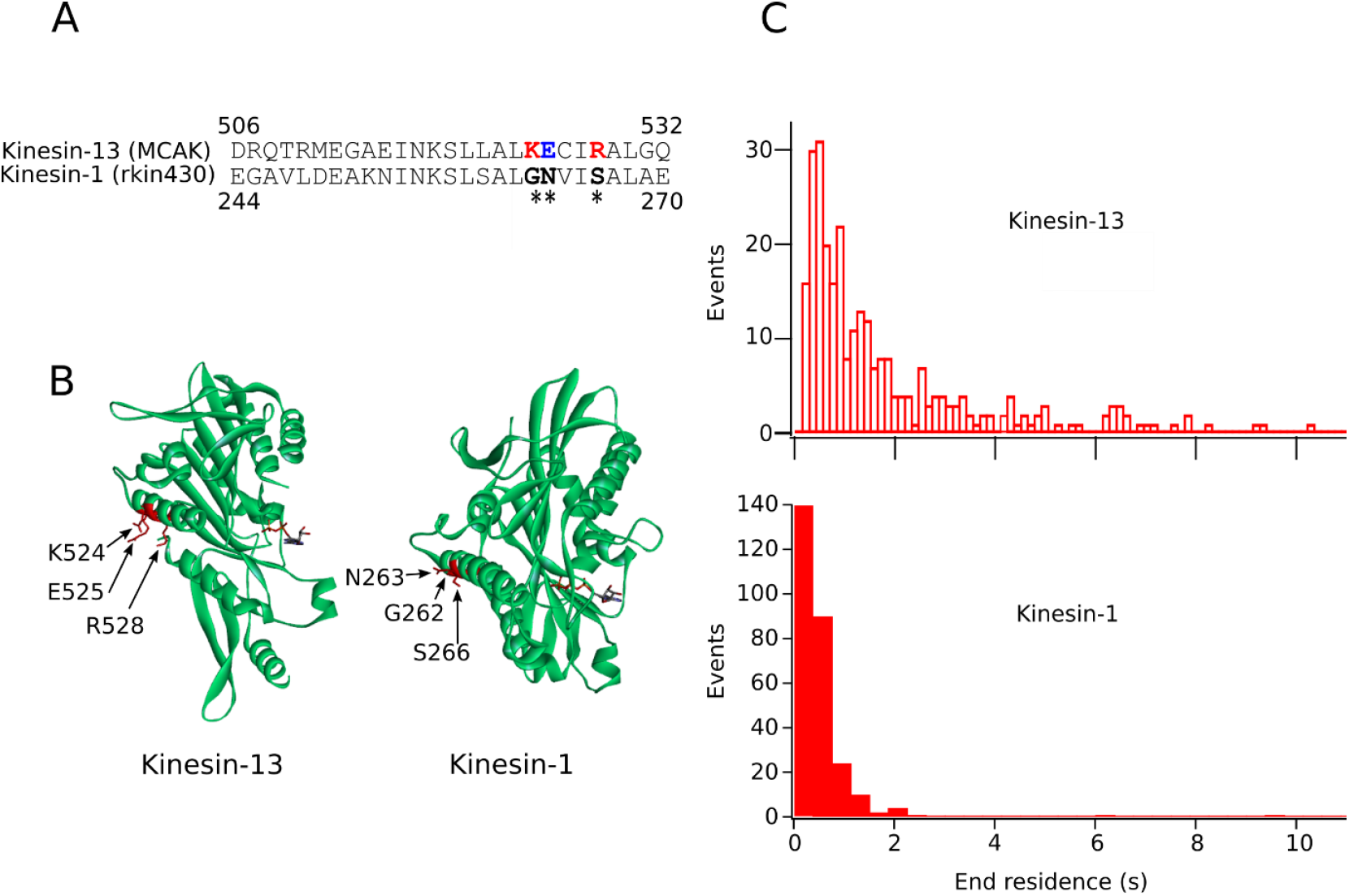
Kinesin 1 and Kinesin 13 show differences in sequence and structure at the α4 helix and behave differently at the microtubule end. A) Alignment of MCAK and rkin430 sequences for the α4 helix – MCAK residues (506-532) and rkin430 residues (244-270). Asterisks indicate the residues we have mutated, which are coloured red for positive, blue for negative and black for neutral. B) Structures of the Kinesin 13 and Kinesin 1 motor domains (MCAK *Homo sapien* PDBID: 2HEH, Kif5B *Homo sapien* PDBID:3J8Y) MCAK residues K524, E525 and R528 and the corresponding residues G262, N263 and S266 in Kif5B are shown as stick structures and highlighted in red. C) Histograms showing the number of interaction events with the microtubule end of a specific duration; MCAK (n=276) and rkin430 (n=273).

### Wild type Kinesin-1 has a short microtubule end residence time relative to Kinesin-13

We expressed and purified the wild type Kinesin-1 truncation, rKin430GFP-h6 as previously described (Rogers, Weiss et al. 2001). We used TIRF microscopy to observe single molecules of rKin430 moving on microtubules and used this data to determine the velocity and run length (Table 1). The values obtained of 810 ± 227 nm/s and 3.06 ± 1.16 μm, respectively, are comparable to values described previously in the literature for this Kinesin-1 construct (Korten and Diez 2008, Ruhnow, Zwicker et al. 2011). We also used this data to determine the microtubule end residence time for a Kinesin-1. Wild type rKin430GFP had a microtubule end residence time of 0.35 ± 0.01 s (Figure 1C), which is in agreement with that previously measured in different buffer conditions of <0.3s (Varga, Leduc et al. 2009). Thus, as expected, the microtubule end residence time for a Kinesin-1 is significantly shorter than the value of 2.03 ± 0.13 s previously measured for the Kinesin-13, MCAK (Figure 1C and (Patel, Belsham et al. 2016)). There is also a clear difference in the distribution of end residence times between the Kinesin-1 and Kinesin-13. Whilst both display many short (<2s) end residence events, for MCAK these make up only 68% of all end residence events, whereas for the Kinesin-1 events shorter than 2s duration make up 97% of the distribution. The Kinesin-13 makes many long interactions with the microtubule end ranging up to ^∼^10s, whilst the Kinesin-1 has few end interaction events longer than ^∼^2s. This contrast in behaviour at the microtubule end is expected as it is a fundamental requirement that a kinesin that regulates microtubule dynamics should be able to recognise the microtubule end, whereas a purely translocating kinesin has no such requirement.

**Table 1:**
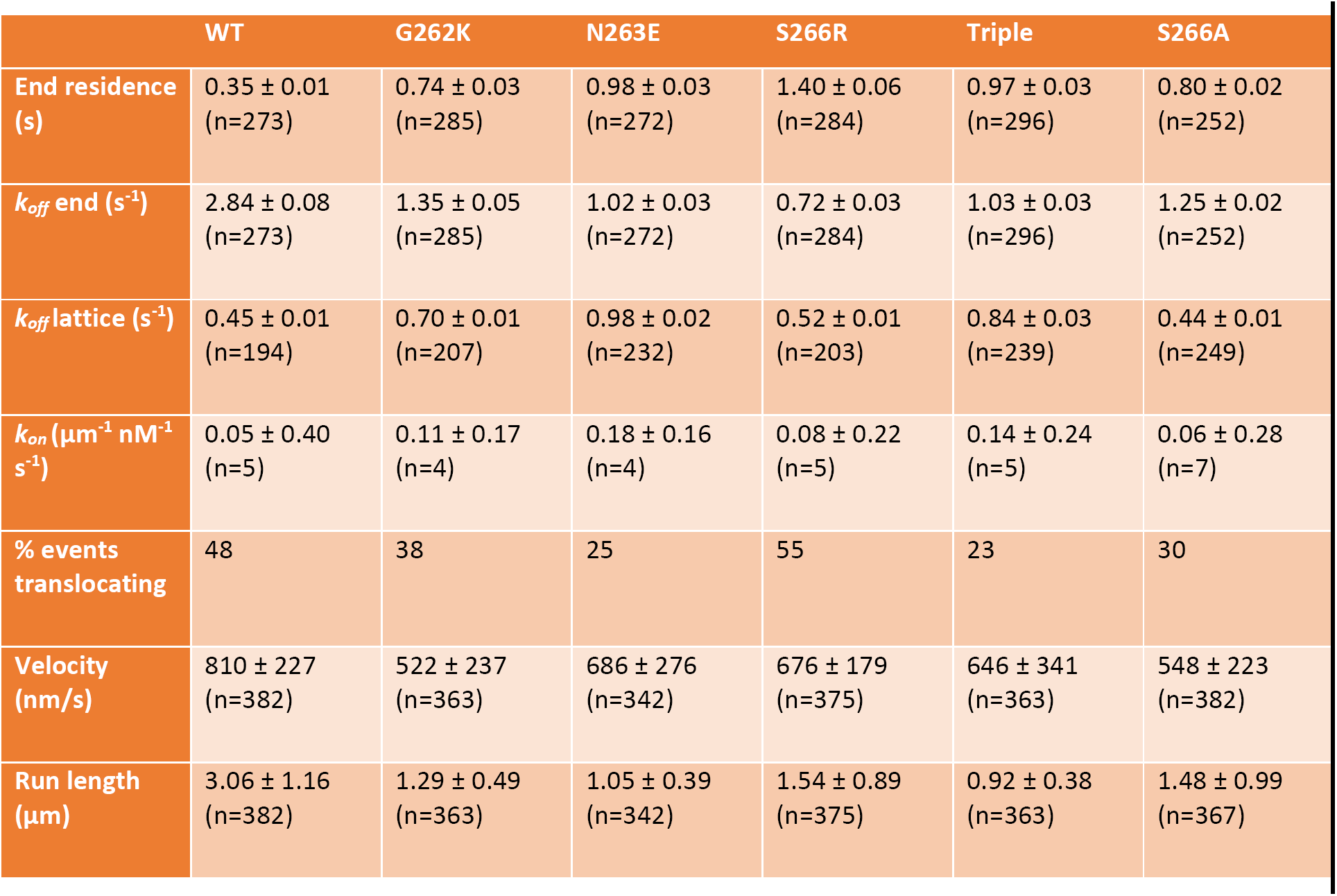
Compiled values for end residence, k_off_, k_on_, translocating percentage, velocity and run length of WT rkin430 and the mutants G262K, N263E, S266R and a triple mutant. Errors are standard deviation. End residence was calculated as the reciprocal of *k*_*off*_. *k*_*off*_ end and lattice were calculated from fitting the cumulative distribution of residence events to an exponential function. *k*_*on*_ was calculated from the total number of events observed on an microtubule, and so here n represents the number of microtubules, rather than the number of events. For *k*_*on*_, velocity and run length mean values are given.

|  | WT | G262K | N263E | S266R | Triple | S266A |
| --- | --- | --- | --- | --- | --- | --- |
| **End residence (s)** | $0.35 \pm 0.01$<br>(n=273) | $0.74 \pm 0.03$<br>(n=285) | $0.98 \pm 0.03$<br>(n=272) | $1.40 \pm 0.06$<br>(n=284) | $0.97 \pm 0.03$<br>(n=296) | $0.80 \pm 0.02$<br>(n=252) |
| **$k_{off}$ end ($s^{-1}$)** | $2.84 \pm 0.08$<br>(n=273) | $1.35 \pm 0.05$<br>(n=285) | $1.02 \pm 0.03$<br>(n=272) | $0.72 \pm 0.03$<br>(n=284) | $1.03 \pm 0.03$<br>(n=296) | $1.25 \pm 0.02$<br>(n=252) |
| **$k_{off}$ lattice ($s^{-1}$)** | $0.45 \pm 0.01$<br>(n=194) | $0.70 \pm 0.01$<br>(n=207) | $0.98 \pm 0.02$<br>(n=232) | $0.52 \pm 0.01$<br>(n=203) | $0.84 \pm 0.03$<br>(n=239) | $0.44 \pm 0.01$<br>(n=249) |
| **$k_{on}$ ($\mu m^{-1} nM^{-1} s^{-1}$)** | $0.05 \pm 0.40$<br>(n=5) | $0.11 \pm 0.17$<br>(n=4) | $0.18 \pm 0.16$<br>(n=4) | $0.08 \pm 0.22$<br>(n=5) | $0.14 \pm 0.24$<br>(n=5) | $0.06 \pm 0.28$<br>(n=7) |
| **% events translocating** | 48 | 38 | 25 | 55 | 23 | 30 |
| **Velocity (nm/s)** | $810 \pm 227$<br>(n=382) | $522 \pm 237$<br>(n=363) | $686 \pm 276$<br>(n=342) | $676 \pm 179$<br>(n=375) | $646 \pm 341$<br>(n=363) | $548 \pm 223$<br>(n=382) |
| **Run length ($\mu m$)** | $3.06 \pm 1.16$<br>(n=382) | $1.29 \pm 0.49$<br>(n=363) | $1.05 \pm 0.39$<br>(n=342) | $1.54 \pm 0.89$<br>(n=375) | $0.92 \pm 0.38$<br>(n=363) | $1.48 \pm 0.99$<br>(n=367) |

### Introducing Kinesin-13 residues into the α4 helix of a Kinesin-1 increases its microtubule end residence time

Our data shows that the motor domain of a Kinesin-13 has an enhanced ability to distinguish the microtubule end from the lattice relative to a Kinesin-1 motor domain. To further understand microtubule end recognition and to determine if it is possible to confer the ability to recognise the microtubule end onto a Kinesin-1, we created three Kinesin-1 mutants, G262K, N263E and S266R, in which three residues critical to MCAK’s ability to distinguish the microtubule end from the lattice were substituted in for the corresponding residues in the α4 helix of rKin430 (Figure 1A). We also created a triple mutant which contained all three point mutations. We expressed and purified these Kinesin-1 variants and used single molecule TIRF microscopy to examine their behaviour on microtubules (Figure 2A). All four Kinesin-1 variants had longer microtubule end residence time than wild type Kinesin-1. The end residence times calculated from the distribution of end interaction times for each variant was 0.74 s, 0.98 s, 1.39 s and 0.97 s for G262K, N263E, S266R and the triple mutant, respectively, compared to 0.35 s for wild type Kinesin-1 (Table 1). The distributions of end interaction events for each variant were confirmed as being statistically significantly different from wild type using the Kolmogorov-Smirnov test (Figure 2 and Table 1). The largest effect on end residence time was observed for the Kinesin-1 mutant S266R. This variant stayed at the microtubule end 4 times longer than wild-type Kinesin-1 and each of the variants had a microtubule end residence time at least 2-fold longer than the wild type (Table 1). The distribution of end residence times for each variant show that the increase in microtubule end residence time is the result of an increased proportion of longer events rather than a general increase in the duration of all events (Figure 2B). In agreement with increased end residence times, the dissociation rate constant (*k*_*off*_) calculated for the microtubule end each variants was decreased with respect to wild type Kinesin-1 (Table 1). By contrast, the microtubule lattice dissociation rate constant was increased for each variant (Table 1 and supplementary Figure S1). This shows that these amino acid substitutions cause a specific increase in end residence time rather than an increased affinity for the microtubule in general. The *k*_*on*_ was not significantly altered from wild-type for any Kinesin-1 variant (Table 1).

**Figure 2:**
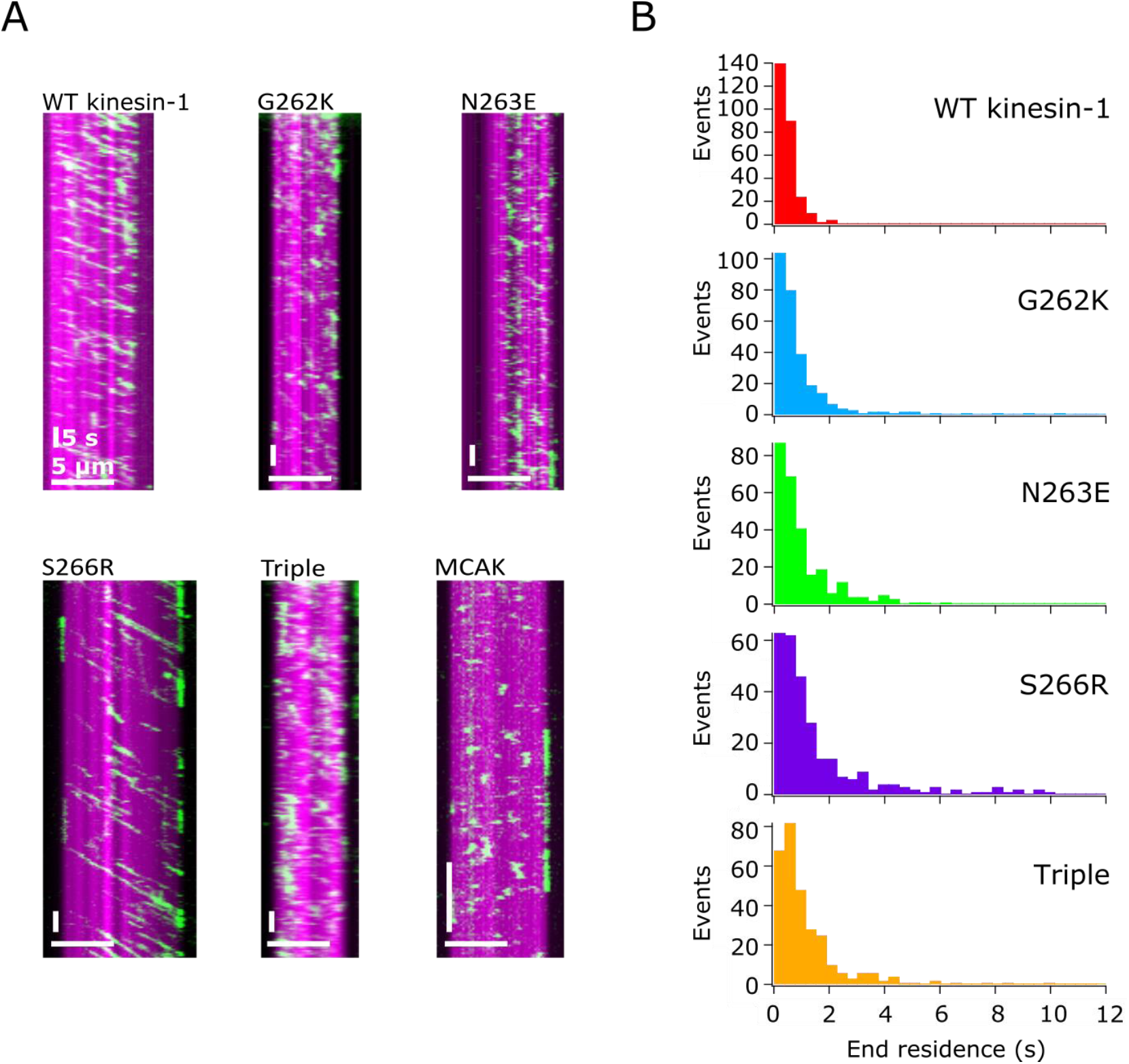
Replacing residues in the α4 helix of Kinesin-1 with those from Kinesin-13 increases the microtubule end residence time. A) Representative kymographs showing the interaction of the wild type Kinesin-1, rKin430-GFP, Kinesin-1 variants and, for comparison, MCAK-GFP (green), with GMPCPP-stabilised, rhodamine labelled microtubules (magenta). B) Histograms showing the number of microtubule end interaction events of a specified duration; WT (n=273), G262K (n=285), N263E (n=272), S266R (n=284), Triple (n=296).

### Increase in end residence time does not correlate with impairment of translocation activity

In addition to the effect on microtubule end residence, the amino acid substitutions studied each caused a reduction in translocation velocity and in run length relative to the wild type Kinesin-1 (Table 1). However, the magnitude of the effect on translocating behaviour does not correlate with the magnitude of the effect on microtubule end residence time. The variant S266R shows the largest increase in end residence but the smallest impact on translocating activity, having the longest run length of all variants tested (Table 1). Further, it can be seen in the kymographs (Figure 2A) that the proportion of molecule that interact with the microtubule in a diffusive manner similar to the Kinesi-13, MCAK, increased for the variants G262K, N263E and the triple mutant relative to wild type Kinesin-1. We observed that 48% of wild type molecules displayed unidirectional translocating activity with 32 % being too short (<750 ms) to allow measurement of velocity and the remaining 20% showing no consistent directional movement. For G262K, N263E and the triple mutant the proportion of translocating events is decreased, with 38, 25 and 23% of events demonstrating unidirectional movement, respectively. By contrast the Kinesin-1 variant that displays the largest increase in end residence time, S266R, does not show any decrease in the proportion of translocating events relative to wild type (55 % translocating).

### Increasing the microtubule end residence of a Kinesin-1 by mutation of the α4 helix is not sufficient to confer microtubule depolymerisation activity

The ability to distinguish between the microtubule end and the lattice is a critical to the function of microtubule depolymerising kinesins (Patel, Belsham et al. 2016). Therefore, to determine whether the increase in microtubule end residence time caused by these amino acid substitution confers microtubule depolymerisation activity to Kinesin-1, we incubated fluorescently labelled GMPCPP-stabilised microtubules with a super-stoichiometric concentration of Kinesin-1, each of the variants and, as a positive control, the Kinesin-13 MCAK in the presence of ATP for 20 minutes. The resulting reaction mixtures were deposited onto cover glasses and imaged by fluorescence microscopy. We observed no significant depolymerisation for wild type Kinesin-1 or any of the Kinesin-1 variants (Figure 3). The appearance of the microtubules after incubation with Kinesin-1 or variants was unchanged, except in the case of S266R which appears to have a microtubule bundling effect (Figure 3). By contrast, under the same conditions microtubules were completely depolymerised by the Kinesin-13, MCAK.

**Figure 3:**
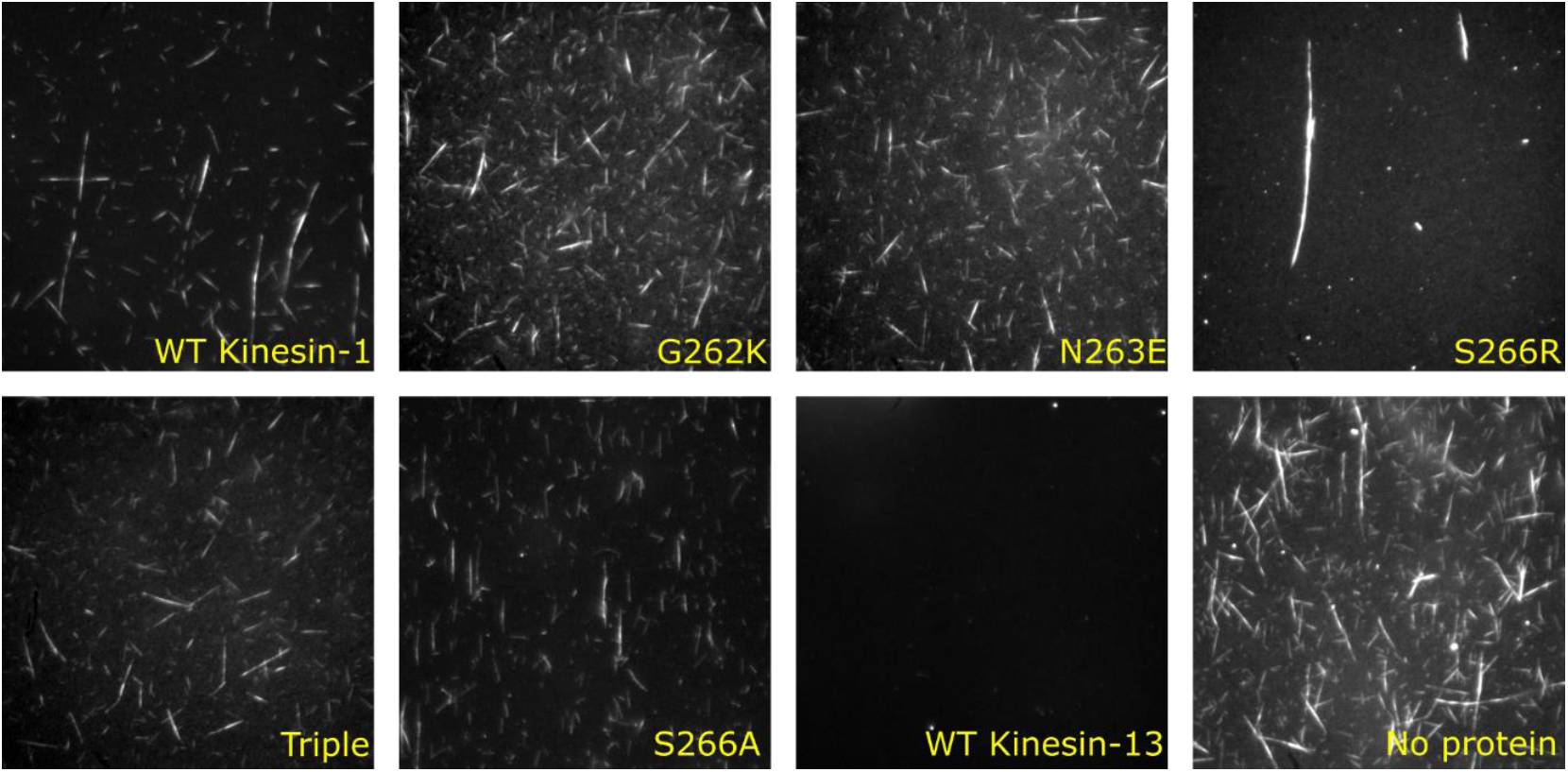
Kinesin-1 mutants with increased microtubule end residence do not depolymerise microtubules. Fluorescence images of the end point of an incubation of GMPCPP-stabilised, rhodamine-labelled microtubules with wild-type Kinesin-1 or the Kinesin-1 mutants G262K, N263E, S266R or the triple mutants containing all three point mutants. The end point of an incubation with wild-type Kinesin-13 (MCAK) and with no addition protein are shown as controls.

### The kinesin motor domain microtubule binding interface is tuned for either lattice or end interactions

To establish whether the increase in end residence time of the Kinesin-1 variants with substituted Kinesin-13 residues was due to introduction of the Kinesin-13 residues or loss of Kinesin-1 residues, we created and analysed the Kinesin-1 variant S266A. Observation of the interaction of this variant with microtubules using single molecule TIRF microscopy, showed that the microtubule end residence time was increased relative to wild type Kinesin-1 (Table 1 and Supplementary Figure S2). However, the end residence time was not increased to the same degree as the introduction of a Kinesin-13 family specific residue at this position (Table 1). The end residence of S266A is 0.8s compared to 1.4s for the Kinesin-13 residue substitution S266R. This suggests that neither lattice binding nor end binding is the default action of the microtubule binding interface of the kinesin motor domain but that the particular amino acid composition found in different kinesin families tunes the affinity for lattice conformation tubulin or end conformation tubulin.

## Discussion

### Introduction of Kinesin-13 family specific resides increases the microtubule end residence time of a Kinesin-1

Despite the high sequence conservation of the kinesin superfamily defining motor domain, kinesins from different families display a remarkable diversity of kinetic and functional properties (Goldstein and Philp 1999, Hirokawa, Noda et al. 2009, Cross and McAinsh 2014, Friel 2018). It is this diversity that allows specialisation of kinesins to particular cellular tasks. A key functional property of kinesins that regulate microtubule dynamics is their ability to recognise the microtubule end. All kinesins which regulate microtubule dynamics studied to date have the ability to distinguish between the microtubule end and the microtubule lattice. The Kinesin-5 Eg5 and the orphan kinesin Kip2, both of which accelerate microtubule polymerization, pause at the microtubule plus end (Chen and Hancock 2015, Hibbel, Bogdanova et al. 2015) The Kinesin-7 (CENP-E) and the non-motile kinesin (NOD), both suggested to promote polymerization, have been shown to localize preferentially to microtubule ends (Cui, Sproul et al. 2005, Sardar, Luczak et al. 2010). The Kinesin-8 (Kip3), which depolymerizes microtubules, resides on the microtubule plus end longer than its stepping time on the microtubule lattice (Varga, Leduc et al. 2009).

The microtubule depolymerising Kinesin-13, MCAK, also has the ability to distinguish the microtubule end from the lattice and ATP turnover by MCAK is maximally stimulated only by microtubule ends (Hunter, Caplow et al. 2003, Friel and Howard 2011). The microtubule end residence time of wild type MCAK is over 4-fold longer than the residence time on the lattice (Patel, Belsham et al. 2016). Mutation of the residues K524, E525 and R528 found in the α4 helix of the microtubule binding face of the motor domain reduce MCAK’s end residence time to match the lattice residence time and impair the microtubule end specific acceleration of ATP turnover (Patel, Belsham et al. 2016). To understand more about microtubule end recognition by kinesins and to determine if microtubule end recognition ability can be conferred to a non-regulating kinesin, we introduced these Kinesin-13 family specific residues, which are critical to microtubule end recognition, into the Kinesin-1 motor domain. We used the Kinesin-1 construct rKin430 (Rogers, Weiss et al. 2001) and showed that its microtubule end residence time was ^∼^6-fold shorter than the Kinesin-13 MCAK, 0.35 s and 2.03 s, respectively (Table 1 and (Patel, Belsham et al. 2016). Thus, confirming that specialist translocating kinesins do not possess the same microtubule end recognition ability characteristic of microtubule regulating kinesins.

Introduction of Kinesin-13 family specific residues into a Kinesin-1 by creating the variants G262K, N263E and S266R increased the end residence time between 2- and 4-fold over the wild type. None of the mutations gave Kinesin-1 the same end residence time as wild type MCAK. Interestingly, the effect of introducing all three mutants to the same construct in a triple mutant is not additive. Rather, the behaviour of the triple mutant is similar to the single mutant N263E suggesting that this position in the sequence is pivotal to the balance between microtubule lattice and microtubule end affinity. The cryo-EM structure of Kinesin-1 in complex with microtubules (PDB ID: 3J8Y, (Shang, Zhou et al. 2014)) shows the N263 side chain is oriented directly towards the tubulin interface, which may explain its dominance in the microtubule interaction. In contrast, the S266 side chain is oriented parallel to the tubulin dimer, which could explain why mutation of this residue has a more pronounced effect on end binding than on translocation activity. This residue may become more important in complex with curved tubulin, as is likely found at the microtubule end, rather than tubulin in a straighter conformation when embedded in the lattice.

### Microtubule end recognition is not sufficient to confer ability to regulate microtubule dynamics

The Kinesin-13 family specific residues we substituted into Kinesin-1 are critical to the ability of MCAK to recognise the microtubule end which promotes ADP dissociation from the nucleotide binding site resulting in the formation of a depolymerisation competent MCAK-tubulin complex. Introduction of these residues to a Kinesin-1 motor domain, however, does not confer depolymerisation activity to Kinesin-1. This suggests that increasing the affinity of the kinesin motor domain for the microtubule end is not sufficient to promote microtubule depolymerisation. It is possible that the end residence time of Kinesin-1 has not been sufficiently increased. However, it is more likely that regions in addition to the α4 helix are crucial to depolymerisation activity. The Kinesin-13 family specific Loop 2 has been shown to be required for microtubule depolymerisation activity (Ogawa, Nitta et al. 2004, Shipley, Hekmat-Nejad et al. 2004, Asenjo, Chatterjee et al. 2013, Wordeman, Decarreau et al. 2016).

### The kinesin motor domain is tuneable to favour either interaction with the microtubule lattice or microtubule end

In addition to their effect in increasing microtubule end residence time, these substitutions to the Kinesin-1 motor domain altered the velocity, run length and proportion of directional events. However, the effect on translocating activity was not necessarily connected to the effect on microtubule end interaction. For all Kinesin-1 variants, along with the end residence time increase, there was a penalty to both the velocity and run length of translocating events. However, the magnitude of the penalty did not correlate with the magnitude of increase in end residence time. The variant which caused the greatest increase in end residence, S266R, had the smallest decrease in run length. S266R also had a similar impact on velocity to N263E and the triple mutant, each of which had only 60% of the effect on end residence caused by the S266R mutation. Further, the Kinesin-1 mutants G262K, N263E and the triple mutant showed an increase in the proportion of diffusive microtubule interaction events rather than directed movement. However, S266R, the variant with the largest increase in end residence time, had no negative impact on the proportion of translocating events. These data indicate that the motor domain-tubulin interface formed on the lattice is subtlety different from that formed at the microtubule end and highlights the potential to tune this interface to favour either lattice or end binding. This has interesting implications for our understanding of the molecular mechanism of kinesins which both translocate and regulate microtubule dynamics and for our ability to manipulate the function of the kinesin motor domain.

## Methods

### Preparation of kinesin

Rat kinesin-1 rKin430-GFP-h6 and all variants were expressed in BL21 *E*. *coli* cells grown in fresh LB media with 0.1 mg/ml ampicillin. Cells were grown overnight and then seeded at a 1:140 dilution, grown up to an OD_595_ of 0.6 and then cooled and grown overnight at 18 ^°^C with 1 mM IPTG. The bacterial pellet was lysed in 50 mM sodium phosphate pH 7.5, 100 mM NaCl, 1 mM MgCl_2_, 10 μM ATP, 5 mM β-mercaptoethanol, with EDTA-free protease inhibitor cocktail tablet (Roche) using a cell disrupter at 35kpsi. After centrifugation at 75,000 x g for 1 hour at 4 ^°^C, the cleared lysate was loaded onto a HisTrap Q column (GE Healthcare). The column was washed with anion buffer (50 mM PIPES pH 6.9, 5 mM MgCl2, 1 mM DTT) with 100 mM NaCl and protein was eluted with anion buffer with 200 mM NaCl. The eluate was then loaded onto a HisTrap HP column (GE Healthcare) and washed with His buffer (50 mM sodium phosphate pH 7.5, 300 mM KCl, 5% glycerol, 1mM MgCl^2^, 10 mM β-mercaptoethanol, 0.1 mM ATP) with 75 mM imidazole, and eluted with His buffer with 300 mM imidazole, 10 % sucrose and 10 mM ATP. Kinesin concentrations are given as monomer concentrations.

### Microtubules

Porcine brain tubulin was purified as described previously (Castoldi and Popov 2003). Fluorescently labelled microtubules were prepared as described previously (Patel, Belsham et al. 2016).

### Single molecule TIRF assays

Single molecules of rkin430-GFP were observed on immobilised, GMPCPP-stabilised, rhodamine labelled, microtubules using TIRF microscopy as described previously(Patel, Belsham et al. 2016), but using BRB12 buffer (12 mM PIPES/KOH pH 6.9, 1 mM EGTA, 1 mM MgCl_2_, 1 mM ATP, 0.1 % Tween 20, 0.1 mg.ml^-1^, BSA, 1 % 2-mercaptoethanol, 40 mM glucose, 40 mg.ml^-1^ glucose oxidase, 16 mg.ml^-1^ catalase). A single image of rhodamine-labelled microtubules was merged with a stack of GFP images, taken every 374 ms (134 ms for MCAK). Kymographs for individual microtubules were used to measure the time individual kinesin molecules spent at the microtubule end and on the lattice. Events were defined by fluorescence above background in 2 or more pixels in the horizontal and 1 or more pixel in the vertical and must be separated by more than 1 non-event pixel in either direction to count as separate events. Translocating events were defined as events which moved in a uni-directional manner through the horizontal and vertical axis simultaneously.

### Depolymerisation assays

GMPCPP-stabilised, rhodamine labelled microtubules were incubated with 40 nM kinesin and 1 mM ATP in BRB12 for 20 minutes. The microtubules were then flowed into a channel made from poly-lysine coated cover glasses and imaged.

## Acknowledgements

We thank Stefan Diez and Corina Braeuer for providing the rKin430-GFP-h6 expression plasmid and sharing advice on expression and purification protocols. TIRF microscopy was carried out in the University of Nottingham, School of Life Sciences Imaging Unit (SLIM) and we thank Chris Gell, in particular, for his assistance. This work was funded by a BBSRC New Investigator award (BB/K006398/1) to C.T.F., the Royal Society and the University of Nottingham.

## Supplementary Information

**Supplementary Figure S1:**
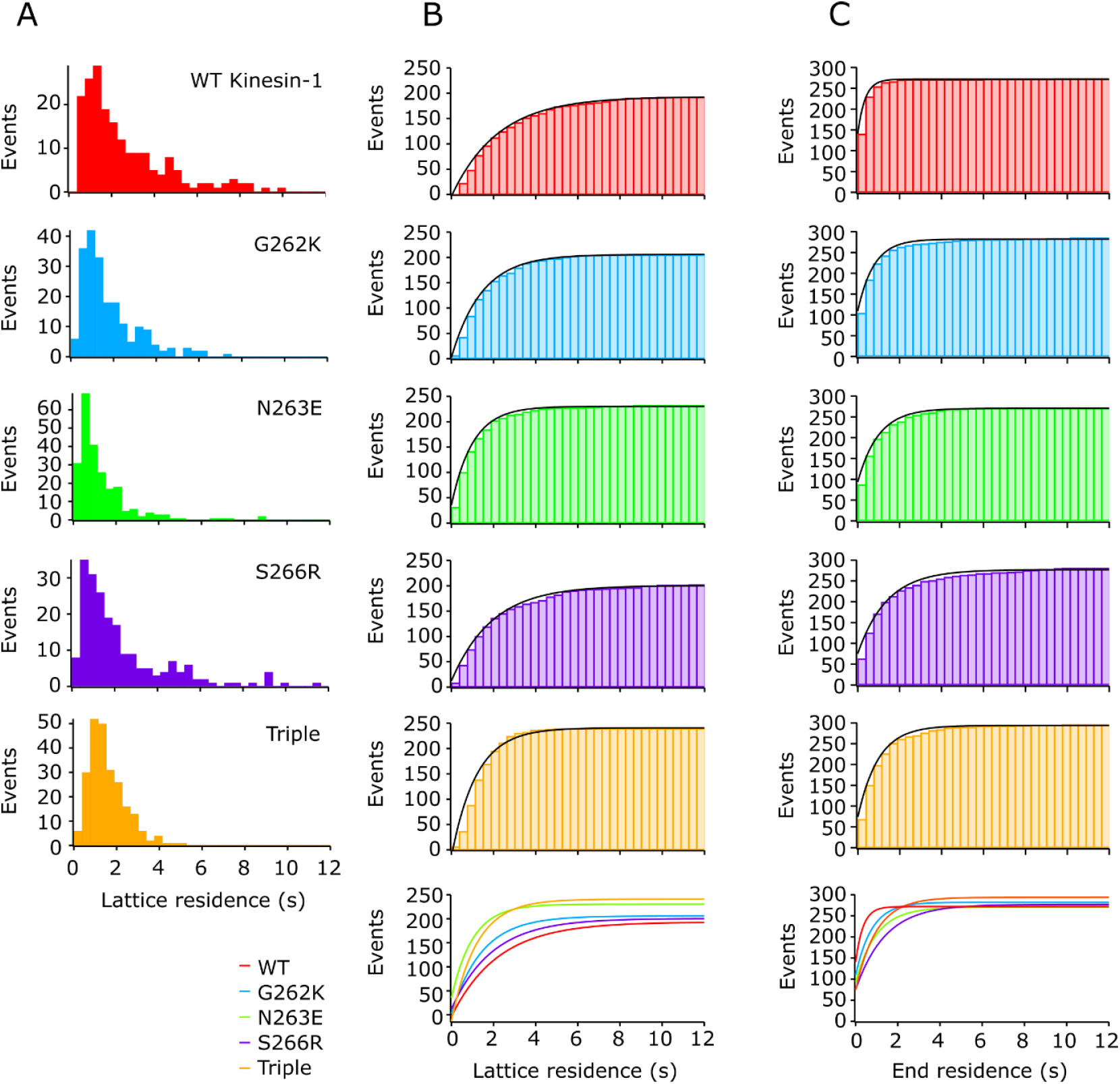
A)Histograms showing the number of lattice residence events of a specified duration: WT (n=194), G262K (n=207), N263E (n=232), S266R (n=203), Triple (n=239), S266A (n=249). B) Cumulative lattice residence histogram with fits shown below. C) Cumulative end residence histogram with fits shown below.

**Supplementary Figure S2:**
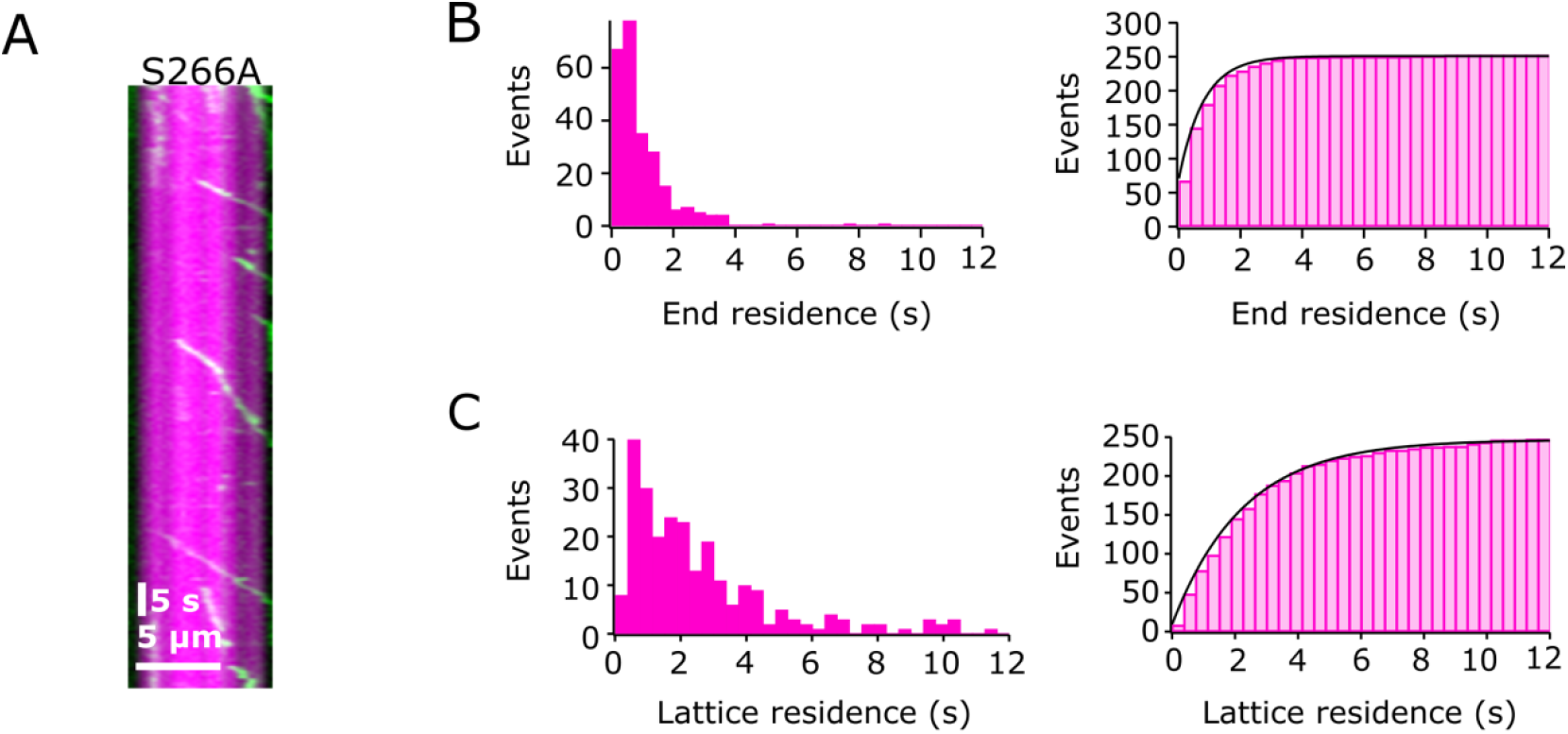
S266A has increased end residence compared to wild type rkin430, but not as much as with the MCAK residues. A) Representative kymographs showing the interaction of GFP-tagged rkin430S266A (green), with GMPCPP-stabilised, rhodamine labelled microtubules (magenta). B) Histogram showing the number of interaction events of S266A with the microtubule end of a specific duration (n=249).

